# ASC oligomer favor caspase-1^CARD^ domain recruitment after intracellular potassium efflux

**DOI:** 10.1101/2020.01.27.921239

**Authors:** Fátima Martín-Sánchez, Vincent Compan, Ana Tapia-Abellán, Ana I. Gómez-Sánchez, María C. Baños, Florian I. Schmidt, Pablo Pelegrín

**Affiliations:** Molecular Inflammation Group, Biomedical Research Institute of Murcia IMIB-Arrixaca, University Clinical Hospital Virgen de la Arrixaca, 30120 Murcia, Spain.; IGF, Univ. Montpellier, CNRS, INSERM, Montpellier, France.; Laboratory of Excellence in Ion Channel Science and Therapeutics (Labex ICST), France.; Institute of Innate Immunity, University of Bonn, Bonn, Germany.; Manchester Collaborative Centre for Inflammation Research, The University of Manchester, Manchester, U.K.; Interfaculty Institute for Cell Biology, Department of Immunology, University of Tübingen, Tübingen, Germany.

**Keywords:** Inflammasome, BRET, NLRP3, nigericin, macrophage, inflammation

## Abstract

Signaling through the inflammasome is important for the inflammatory response. Low concentrations of intracellular K^+^ are associated with the specific oligomerization and activation of the NLRP3 inflammasome, a type of inflammasome involved in sterile inflammation. Subsequent to NLRP3 oligomerization, ASC protein binds and form oligomeric filaments culminating in large protein complexes named ASC specks. ASC specks are also initiated from different inflammasome scaffolds, as AIM2, NLRC4 or Pyrin. ASC oligomers induce the recruitment of caspase-1 through interactions between their respective caspase activation and recruitment domains (CARD), and favoring its activation. So far ASC oligomerization and caspase-1 activation are considered as a K^+^-independent process. Here we found that ASC oligomers change their structure upon low intracellular K^+^ independently of NLRP3 and allow the ASC^CARD^ domain to be more accessible for the recruitment of pro-caspase-1^CARD^ domain. Therefore, conditions that decrease intracellular K^+^ not only drive NLRP3 responses, but also enhance the recruitment of pro-caspase-1 by ASC specks formed by different inflammasomes, indicating that intracellular K^+^ homeostasis is a key regulatory step for inflammasome regulation.

## Introduction

The nucleotide-binding domain and leucine-rich repeat-containing receptor with a pyrin domain 3 (NLRP3) inflammasome is a regulator of inflammation and immunity. NLRP3 is a multiprotein complex, whose oligomerization occurs in response to multiple pathogen and non-infectious triggers, including situations where intracellular ion homeostasis is disturbed [1, 2]. Compounds that induce a decrease in intracellular K^+^ concentration, as P2X7 receptor activation, TWIK2 ion channel opening, cell swelling, pore-forming toxins or selective K^+^-ionophores, all lead to NLRP3 activation [2–7]. Although the mechanism behind NLRP3 oligomerization in response to K^+^-efflux is not well understood, recently it has been suggested that lowering intracellular K^+^ leads to Golgi rearrangement and NLRP3 binding to negatively charged lipids on the Golgi to favor its activation and later interaction with never in mitosis A-related kinase 7 (NEK7) [8–10]. NLRP3 oligomer binds the adaptor ASC (apoptosis-associated speck-like protein with a caspase recruitment domain) and promote ASC oligomerization in filaments that form large protein complexes called ASC specks. ASC oligomerization from NLRP3 oligomers occurs via their pyrin domain (PYD) and recruitment of new ASC subunits into the ASC filament continue via PYD-PYD interactions [11–13]. In these filaments, the ASC caspase recruitment domain (CARD) is exposed to the external side and favor filament binding and recruitment of pro-caspase-1 via CARD-CARD interactions to activate caspase-1 within inflammasome complexes [13–16]. Active caspase-1 is now able to process pro-inflammatory cytokines of the interleukin (IL)-1 family and also induces their release via the processing of gasdermin D and formation of plasma membrane pores [17]. Uncontrolled gasdermin D plasma membrane pores will lead to a specific type of cell death termed pyroptosis, that will amplify the inflammatory response by releasing intracellular content, including inflammasome oligomers [17–19]. Besides the NLRP3 inflammasome, ASC could also oligomerize and activate caspase-1 in response to other inflammasome scaffolds, as the AIM2, Pyrin or NLRC4 [20–22]. The oligomerization of these inflammasomes is independent of the concentration of intracellular K^+^, and sensors are rather promoted by the presence of cytosolic nucleic acids, flagellin or type III secretion system proteins, or RhoA GTPase inhibition [20,21,23,24]. So far, the oligomerization of ASC has been thought to be independent of intracellular K^+^ concentration. Here, we describe that the ASC speck could change its structure in conditions of low intracellular K^+^ to allow ASC^CARD^ domains to be more accessible and enhance the recruitment of pro-caspase-1^CARD^ domains. Our results show that the activity of ASC oligomerized by inflammasome sensors other than NLRP3 can be enhanced by a decrease of intracellular K^+^. These conditions could occur during the initial phases of gasdermin D plasma membrane permeabilization and would amplify caspase-1 activation.

## Results

### NLRP3, but not ASC, oligomerize in response to K^+^ efflux

The expression of the inflammasome proteins NLRP3, ASC and pro-caspase-1 in HEK293 cells result in functional reconstitution of the NLRP3 inflammasome able to activate caspase-1 in response to intracellular K^+^ decrease induced by hypotonic cell swelling (**Fig. 1A**), confirming that HEK293 cells are suitable platforms to study NLRP3 inflammasome activation [3,8,25,26]. NLRP3 and ASC expression in HEK293 results in the co-oligomerization of these proteins only after nigericin or hypotonic stimulation (**Fig. 1B**). As expected, the oligomerization of NLRP3 was due to K^+^ efflux, since it was blocked when nigericin was added in a high K^+^ buffer (**Fig. S1A,B**). Also, as previously described [19, 27], the pathological mutation of NLRP3 D303N associated with cryopyrin associated periodic syndromes (CAPS) resulted in spontaneous oligomerization of NLRP3 when expressed in HEK293 cells without any stimulation (**Fig. 1C**). This oligomerization was not affected by K^+^ efflux, as nigericin or hypotonic stimulation did not affect basal NLRP3 D303N oligomers (**Fig. S1C**). When ASC is expressed with the D303N mutated NLRP3, ASC assembly led to the convergence of NLRP3 oligomers into single structures per cell (**Fig. 1C**).

**Figure 1.**
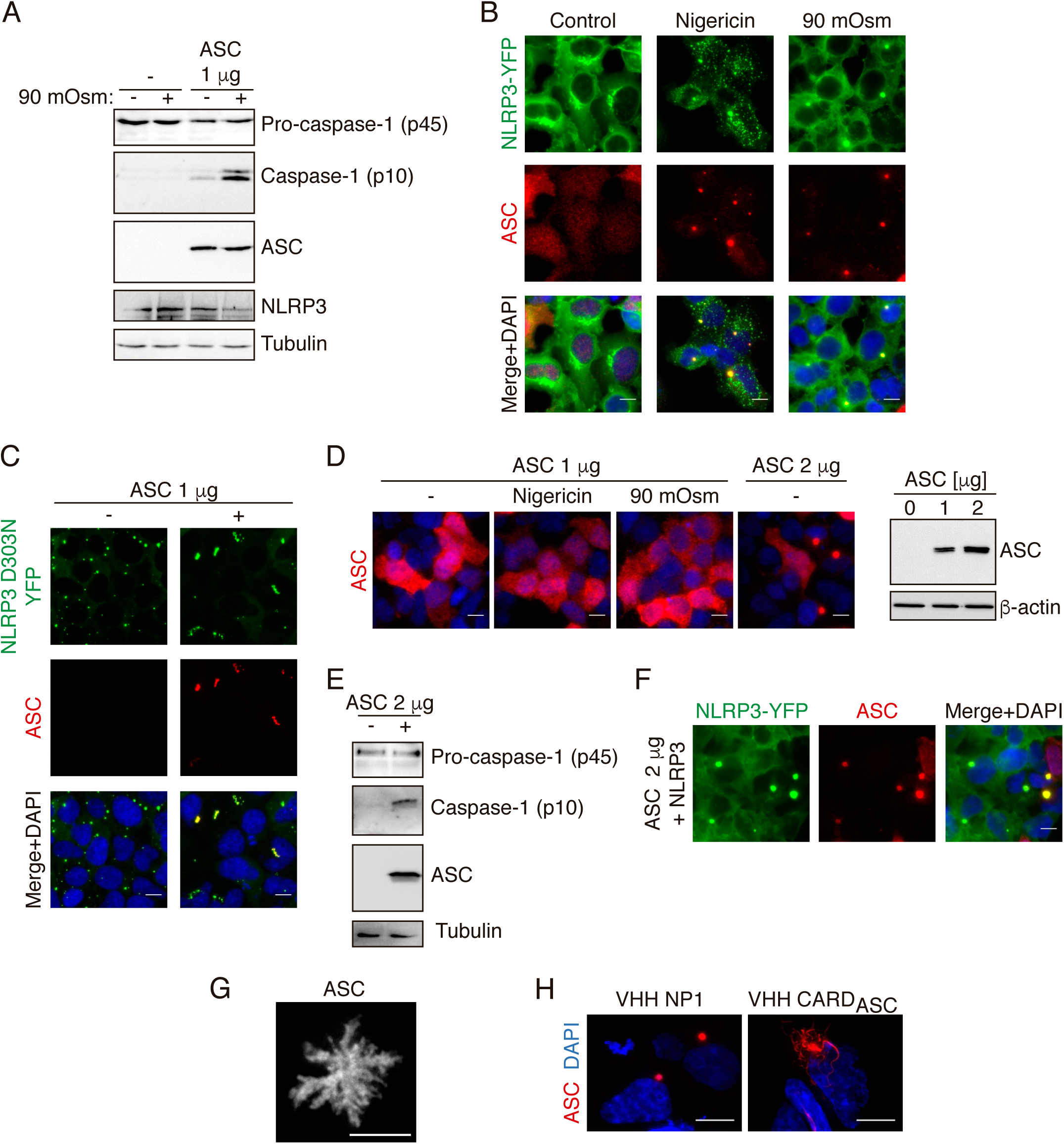
K^+^ efflux induces NLRP3, but not ASC, oligomerization. **(A)** Caspase-1, ASC, NLRP3 and tubulin immunoblot of HEK293 cells transfected with plasmids encoding for pro-caspase-1, NLRP3 and in the last two lanes with ASC (1 μg). After 16 h of transfection cells where exposed to hypotonic buffer (90 mOsm) for 1 h as indicated before cell lysis. **(B)** Representative fluorescent photomicrographs of NLRP3-YFP HEK293 stable cell line transfected with a plasmid encoding for ASC (1 μg) and unstimulated (control) or stimulated with either nigericin (10 μM, 30 min) or hypotonic solution (90 mOsm, 1 h) as indicated. NLRP3-YFP was revealed by direct excitation of YFP (green), ASC was stained with polyclonal α-PYD_ASC_ antibody (AF647, red) and nuclei were revealed with DAPI (blue); scale bar 10 μm. **(C)** Representative fluorescent photomicrographs of NLRP3-YFP D303N HEK293 stable cell line transfected or not with a plasmid encoding for ASC (1 μg) as indicated. NLRP3-YFP D303N was revealed by direct excitation of YFP (green), ASC was stained with polyclonal α-PYD_ASC_ antibody (AF647, red) and nuclei were revealed with DAPI (blue); scale bar 10 μm. **(D)** Representative fluorescent photomicrographs of HEK293 cells transfected with a plasmid encoding for ASC (1 or 2 μg, as indicated). ASC was stained with polyclonal α-PYD_ASC_ antibody (AF647, red) and nuclei were revealed with DAPI (blue); scale bar 10 μm (left panels). ASC and β-actin immunoblot of cells extracts (right). **(E)** Caspase-1, ASC and tubulin immunoblot of HEK293 cells transfected with plasmids encoding for pro-caspase-1 and in ASC (2 μg). **(F)** Representative fluorescent photomicrographs of NLRP3-YFP HEK293 stable cell line transfected with a plasmid encoding for ASC (2 μg). NLRP3-YFP was revealed by direct excitation of YFP (green), ASC was stained with polyclonal α-PYD_ASC_ antibody (AF647, red) and nuclei were revealed with DAPI (blue); scale bar 10 μm. **(G)** Deconvolved representative fluorescent photomicrographs of HEK293 cells transfected with a plasmid encoding for ASC (2 μg). ASC was stained with monoclonal α-CARD_ASC_ antibody (AF647, red); scale bar 10 μm. **(H)** Deconvolved representative fluorescent photomicrographs of HEK293 cells transfected with plasmids encoding for ASC-RFP (0.1 μg) and for a control nanobody (VHH NP1) or for a nanobody α-CARD_ASC_ (VHH CARD_ASC_). ASC-RFP was revealed by direct excitation of RFP (red) and nuclei were revealed with DAPI (blue); scale bar 10 μm.

The expression of low amounts of ASC alone in HEK293 was unable to trigger oligomerization in response to nigericin or hypotonic solution (**Fig. 1D**), and NLRP3-deficient macrophages expressing ASC do not release IL-1β after nigericin stimulation (**Fig. S1D**), confirming that NLRP3 is the sensor protein responsible for oligomerization of functional inflammasomes in response to K^+^ efflux [7]. However, for higher expression levels of ASC alone, spontaneous oligomerization of ASC was observed in HEK293 (**Fig. 1D**). These oligomers were functional as a platform inducing self-processing of caspase-1 (**Fig. 1E**) and they recruit wild type NLRP3 into the ASC specks without any cell stimulation that decrease intracellular K^+^ (**Fig. 1F**). Therefore, ASC specks formed by high expression levels of ASC present a functional structure. To study the structure of these ASC oligomers, we did high resolution imaging and found ASC oligomers with filamentous nature (**Fig. 1G**). When ASC was expressed together with a nanobody binding to the ASC^CARD^ domain and impairing ASC^CARD^ homotypic interactions [13], ASC oligomerization was found to result in filaments (**Fig. 1H**).

### BRET assay reveals a structural change of ASC specks in response to K^+^ efflux

Spontaneous ASC speck formation induced by high expression was not affected by tagging ASC with either YFP or Luc at C-terminus (**Fig. S2A,B**). Co-expression of ASC-YFP and ASC-Luc results in ASC-specks containing both types of tagged ASC (**Fig. 2A**). BRET signal between ASC-Luc and ASC-YFP inside the specks shows a specific BRET signal (**Fig. 2B,C**). After nigericin or hypotonic stimulation, ASC speck BRET signal decreased (**Fig. 2D**) and this decrease was blocked by high extracellular K^+^ concentration (**Fig. 2E, S2C**), suggesting that the ASC speck, independently of NLRP3, is able to undergo a structural change due to K^+^ efflux. Co-expression of wild-type NLRP3 or D303N mutant increased ASC speck BRET signal (**Fig. 2F**), but did not affect the general profile of the ASC speck structural changes induced by K^+^ efflux (**Fig. 2G**). However, the presence of NLRP3 resulted in a faster and more pronounced decrease of the BRET signal after nigericin or hypotonic stimulation (**Fig. 2G**).

**Figure 2.**
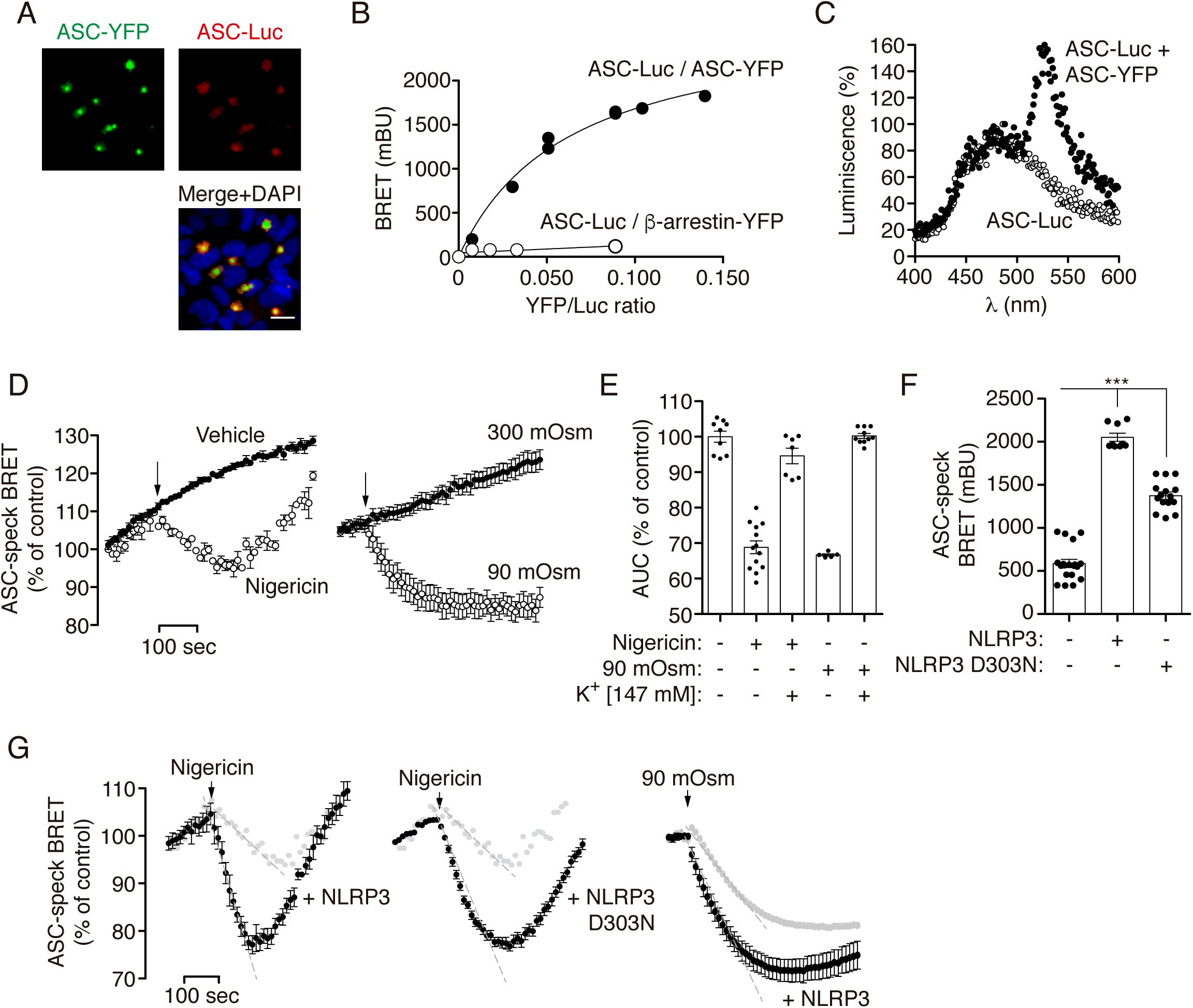
BRET assays reveal a structural change of ASC specks in response to K^+^ efflux. **(A)** Representative fluorescent phtomicrographs of HEK293 cells transfected with plasmids encoding for ASC-YFP (0.2 μg) and ASC-Luc (0.1 μg). ASC-YFP was revealed by direct excitation of YFP (green), ASC-Luc was stained with α-Luc antibody (AF647, red) and nuclei were revealed with DAPI (blue); scale bar 10 μm. **(B)** BRET saturation curves for HEK293 cells transfected with a constant concentration of ASC-Luc and increasing amounts of the BRET acceptor ASC-YFP (black circles) or β-arrestin-YFP (white circles). mBU, milliBRET units. **(C)** Emission spectra of HEK293 cells expressing ASC-Luc (white circles) or ASC-Luc with ASC-YFP (black circles) after adding coelenterazine h. Note the peak emission at 535 nm as the BRET among ASC-Luc and ASC-YFP inside the ASC speck. **(D)** Kinetic of net BRET signal in HEK293 cells transfected with the BRET donor ASC-Luc and the acceptor ASC-YFP in response to nigericin (left) or hypotonic solution (90 mOsm, right). **(E)** Area under the curve (AUC) of the net BRET signal kinetic in HEK293 cells transfected with the BRET donor ASC-Luc and the acceptor ASC-YFP in response to nigericin or hypotonic solution (90 mOsm) recorded in a buffer containing 147 mM of KCl. **(F)** BRET signal in HEK293 cells transfected with the BRET donor ASC-Luc and the acceptor ASC-YFP in the presence of NLRP3 wild-type or D303N as indicated. **(G)** Kinetic of net BRET signal in HEK293 cells transfected with the BRET donor ASC-Luc and the acceptor ASC-YFP in the presence of NLRP3 wild-type or D303N as indicated, in response to nigericin (left two panels) or hypotonic solution (90 mOsm, right panel); light grey dots represent the BRET signal for ASC-Luc and ASC-YFP in the absence of NLRP3 expression.

### K^+^ efflux induced ASC speck structural changes favor ASC labelling with antibodies

The structural change of the ASC speck due to K^+^ efflux was also evidenced by immunofluorescence experiments and ASC staining with different antibodies. Using anti-Luc antibodies, staining of ASC speck formed by ASC-YFP and ASC-Luc was more intense after nigericin or hypotonic treatment (**Fig. 3A,B**), whereas ASC-YFP fluorescence remained constant whatever the treatment. Change in ASC-Luc staining was transient after nigericin treatment and stable after hypotonic stimulation (**Fig. 3A,B**). The increase in ASC staining observed after nigericin treatment was blocked by high K^+^ extracellular solution (**Fig. 3C**). These data are consistent with the BRET recording kinetics described above (**Fig. 2D**). Similarly, using an anti-ASC antibody against the CARD domain, we observed changes in ASC staining in HEK293 cells overexpressing untagged mouse and human ASC upon hypotonic or nigericin stimulation (**Fig. 3D,E**). This antibody against the ASC^CARD^ domain specifically recognized oligomeric ASC but did not detect soluble ASC neither overexpressed in HEK293 cells or endogenously expressed by mouse macrophages (**Fig. S3A,B**). These data suggest that upon K^+^ efflux, the ASC^CARD^ domain is more accessible to antibody staining.

**Figure 3.**
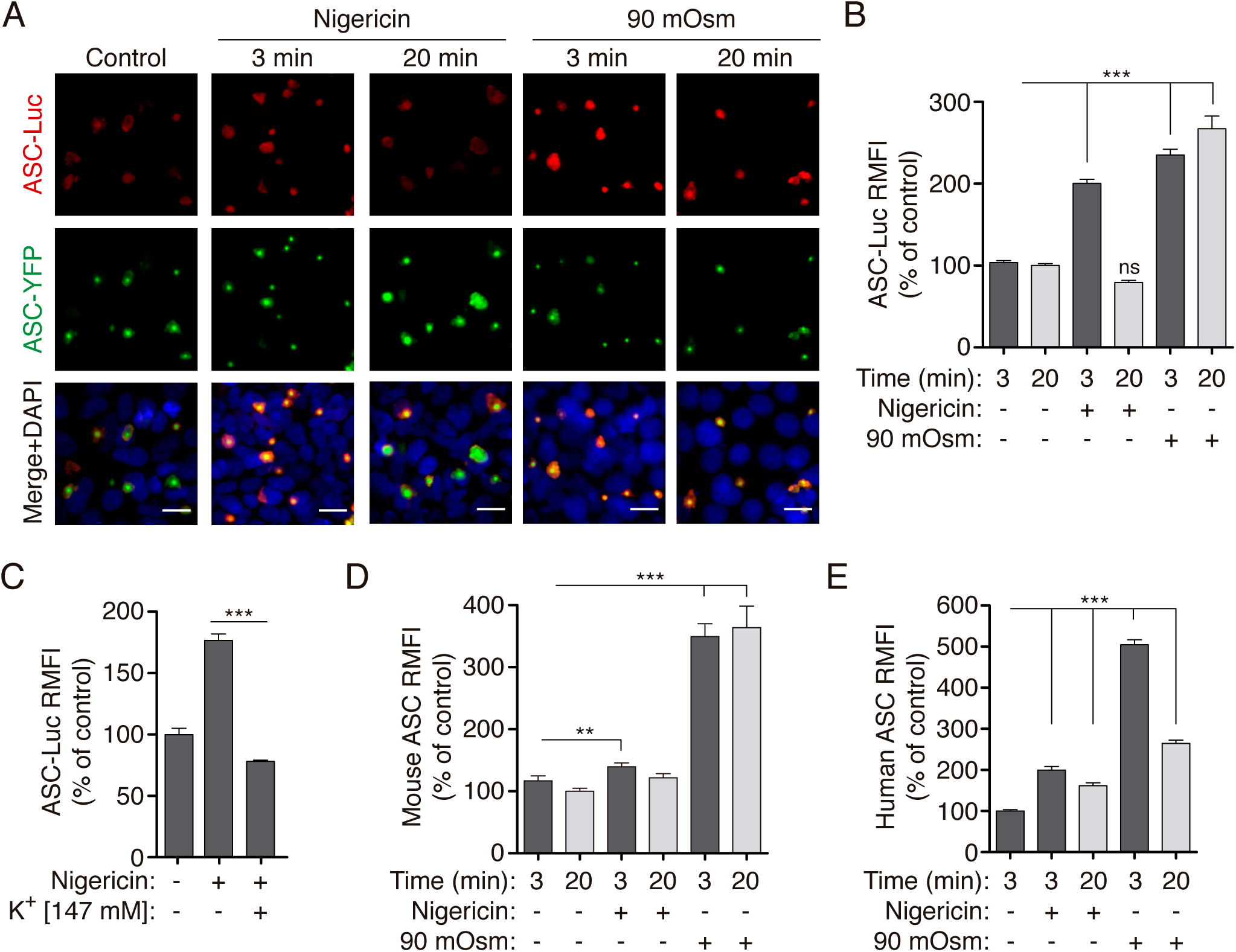
ASC speck structural change in response to K^+^ efflux is revealed with immunofluorescence. **(A)** Representative fluorescent phtomicrographs of HEK293 cells transfected with plasmids encoding for ASC-YFP (0.2 μg) and ASC-Luc (0.1 μg), unstimulated (control) or stimulated with nigericin (10 μM) or hypotonic solution (90 mOsm) for the indicated time. ASC-YFP (green), ASC-Luc was stained with α-Luc antibody (AF647, red) and nuclei were revealed with DAPI (blue); scale bar 10 μm. **(B)** Relative mean fluorescence intensity (RMFI) of ASC-Luc oligomers from HEK293 transfected and stimulated as in (A). Data from *n*= 3 independent experiments and quantification of a total of 2,084 ASC-specks. **(C)** RMFI of ASC-Luc oligomers from HEK293 transfected and stimulated as in (A) but when indicated in the presence of a buffer containing 147 mM KCl. Data from *n*= 3 independent experiments and quantification of a total of 1,278 ASC-specks. **(D,E)** RMFI of ASC oligomers from HEK293 transfected with a plasmid encoding for untagged mouse ASC (2 μg, D) or untagged human ASC (1 μg, D) and stimulated as in (A). ASC was labeled with monoclonal α-CARD_ASC_. Data from *n*= 2-3 independent experiments and quantification of a total of 1,323 (D) and 1,331 (E) ASC-specks.

Without any stimulation, the antibody against the ASC^CARD^ was unable to recognize soluble or oligomeric mouse ASC tagged at Ct with YFP (**Fig. 4A, S3C**). However, after nigericin or hypotonic treatment, ASC-YFP specks were stained (**Fig. 4A,B**) and this staining was completely abolished by high extracellular concentration of K^+^ (**Fig. S3D**). In contrast to staining of the CARD domain, an antibody against the ASC^PYD^ domain was able to detect soluble and oligomeric ASC either untagged (**Fig. 1D**) or tagged with YFP at Ct (**Fig. S3A**) and this staining was not affected by change in intracellular K^+^ concentration (**Fig. S3E**). These data suggest that K^+^ efflux induces a change of ASC speck conformation allowing the ASC^CARD^ domain, but not the ASC^PYD^ domain, to be more accessible to antibody staining.

**Figure 4.**
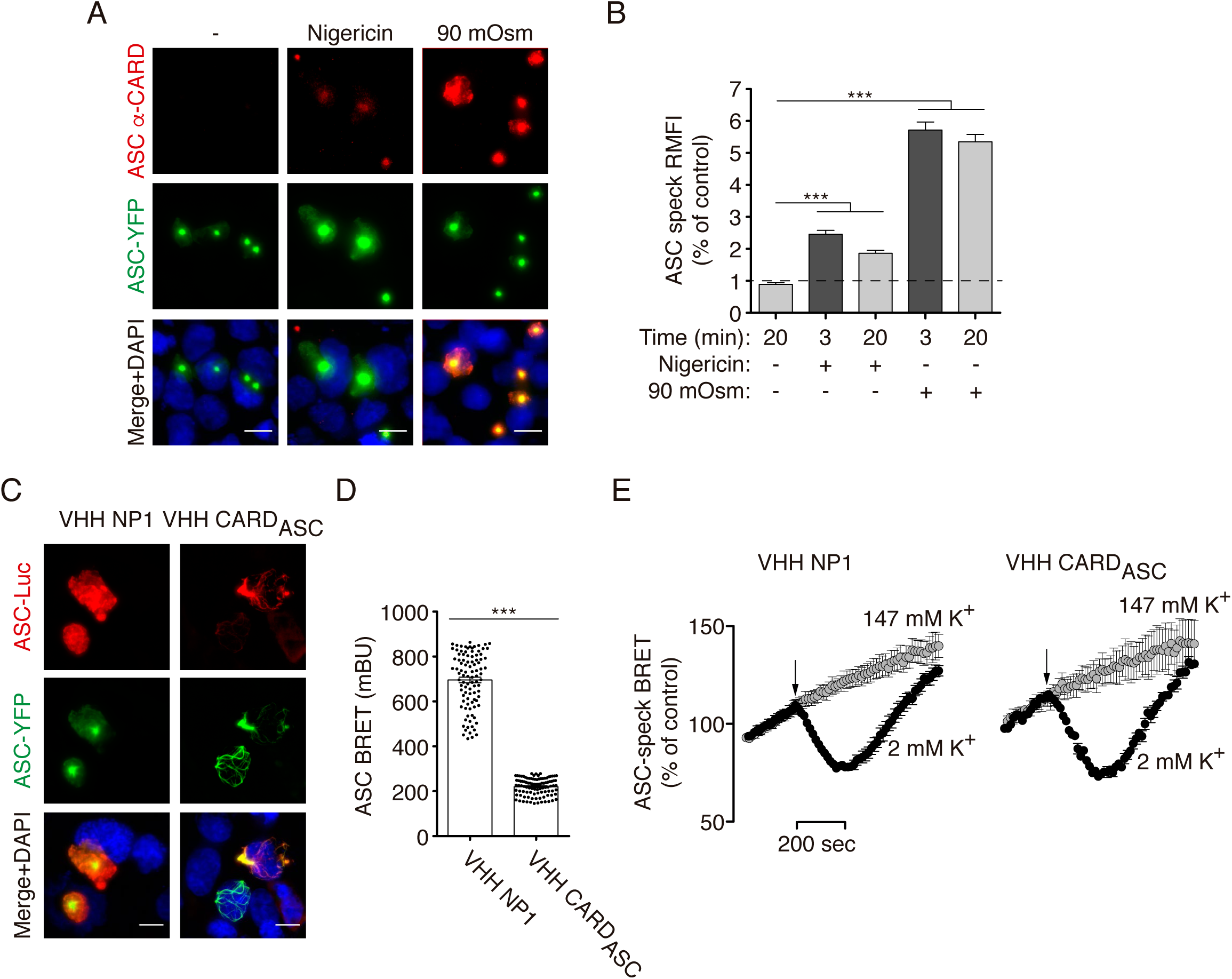
ASC speck structural change is differentially labelled with antibodies. **(A)** Representative fluorescent micrographs of HEK293 cells transfected with a plasmid encoding for ASC-YFP (0.2 μg), unstimulated (control) or stimulated with nigericin (10 μM) or hypotonic solution (90 mOsm) for 3 min. ASC was detected by the YFP fluorescence (green) and stained with monoclonal α-CARD_ASC_ antibody (AF647, red), nuclei were revealed with DAPI (blue); scale bar 10 μm. **(B)** Relative mean fluorescence intensity (RMFI) of ASC-YFP oligomers stained with monoclonal α-CARD_ASC_ antibody from HEK293 transfected and stimulated as in (A). Data from *n*= 2 independent experiments and quantification of a total of 1,103 ASC-specks. **(C)** Representative fluorescent photomicrographs of HEK293 cells transfected with plasmids encoding for ASC-YFP (0.2 μg), ASC-Luc (0.1 μg), and for either control nanobody (VHH NP1, 0.5 μg) or α-CARD_ASC_ nanobody (VHH CARD_ASC_, 0.5 μg). ASC-YFP was revealed by direct excitation of YFP (green), ASC-Luc was stained with α-Luc antibody (AF647, red) and nuclei were revealed with DAPI (blue); scale bar 10 μm. **(D)** BRET signal in HEK293 cells transfected as in (C). Data from *n*= 4 independent experiments. **(E)** Kinetic of net BRET signal in HEK293 cells transfected as in (C) in response to nigericin in normal extracellular K^+^ buffer (2 mM K^+^) or high K^+^ buffer (147 mM K^+^). Data from *n*= 4 independent experiments.

To identify if the change observed in the structure of the ASC speck was due to changes on the PYD-PYD oligomerization filament or the CARD-CARD interactions of different filaments, we used a nanobody binding to the ASC^CARD^ domain and impairing ASC^CARD^ homotypic interactions [13]. with ASC-YFP and ASC-Luc BRET sensors. This resulted in ASC filamentous structures containing both types of ASC (**Fig. 4C**). BRET recording show that ASC filaments presented a lower BRET signal when compared to the BRET signal from the ASC speck co-expressed with a control nanobody (**Fig. 4D**), suggesting that oligomerization of ASC into a speck increased BRET signal over the BRET signal of the ASC filament. After nigericin treatment, we found a similar change in the BRET signal that was abrogated by using a high extracellular K^+^ solution (**Fig. 4E**), suggesting that the structural change in the ASC filament is similar than of the full ASC speck. This supports that the oligomerization of ASC by the ASC^PYD^ domain undergoes structural quaternary changes after K^+^ efflux, allowing the ASC^CARD^ domain to be more accessible.

### K^+^ efflux induced structural change of ASC specks recruit more pro-caspase-1^CARD^

Our results lead us to hypothesize that the change of the ASC speck structure leads to an exposure of the ASC^CARD^ domain in an intracellular environment with low K^+^, possible resulting in a better recruitment of pro-caspase-1. To address this question, we stained ASC specks with cytosolic extracts containing either soluble pro-caspase-1^CARD^ domain tagged with EGFP (unpublished, F.I. Schmidt lab), which would be recruited by ASC^CARD^ interactions, or soluble ASC-RFP that could be mainly recruited by ASC^PYD^ interactions, but also by ASC^CARD^ interactions. By detecting GFP and RFP fluorescence, we found that both the pro-caspase-1^CARD^ domain and ASC-RFP respectively were recruited by already formed ASC specks (**Fig. 5A**). However, ASC antibody against the ASC^CARD^ domain failed to stain ASC-RFP recruited to ASC speck (**Fig. 5B**), since this antibody does not recognize mouse ASC tagged on the Ct in the absence of CARD domain exposure (**Fig. S3A**). Upon K^+^ efflux induced by hypotonic stimulation of HEK293 cells expressing untagged ASC specks, we found an increase in the recruitment of the pro-caspase-1^CARD^ domain, but not of additional ASC-RFP subunits (**Fig. 5C,D**). Staining of ASC with the anti-ASC^CARD^ domain antibody recognize a structural change in the ASC speck upon hypotonic stimulation that was further increased by the recruitment of ASC-RFP into the speck (**Fig. 5E**). This suggests that the recruitment of new proteins via the ASC^CARD^ domain is facilitated in situations of low intracellular K^+^, but the recruitment of new subunits of ASC, possibly via ASC^PYD^ domain, would not.

**Figure 5.**
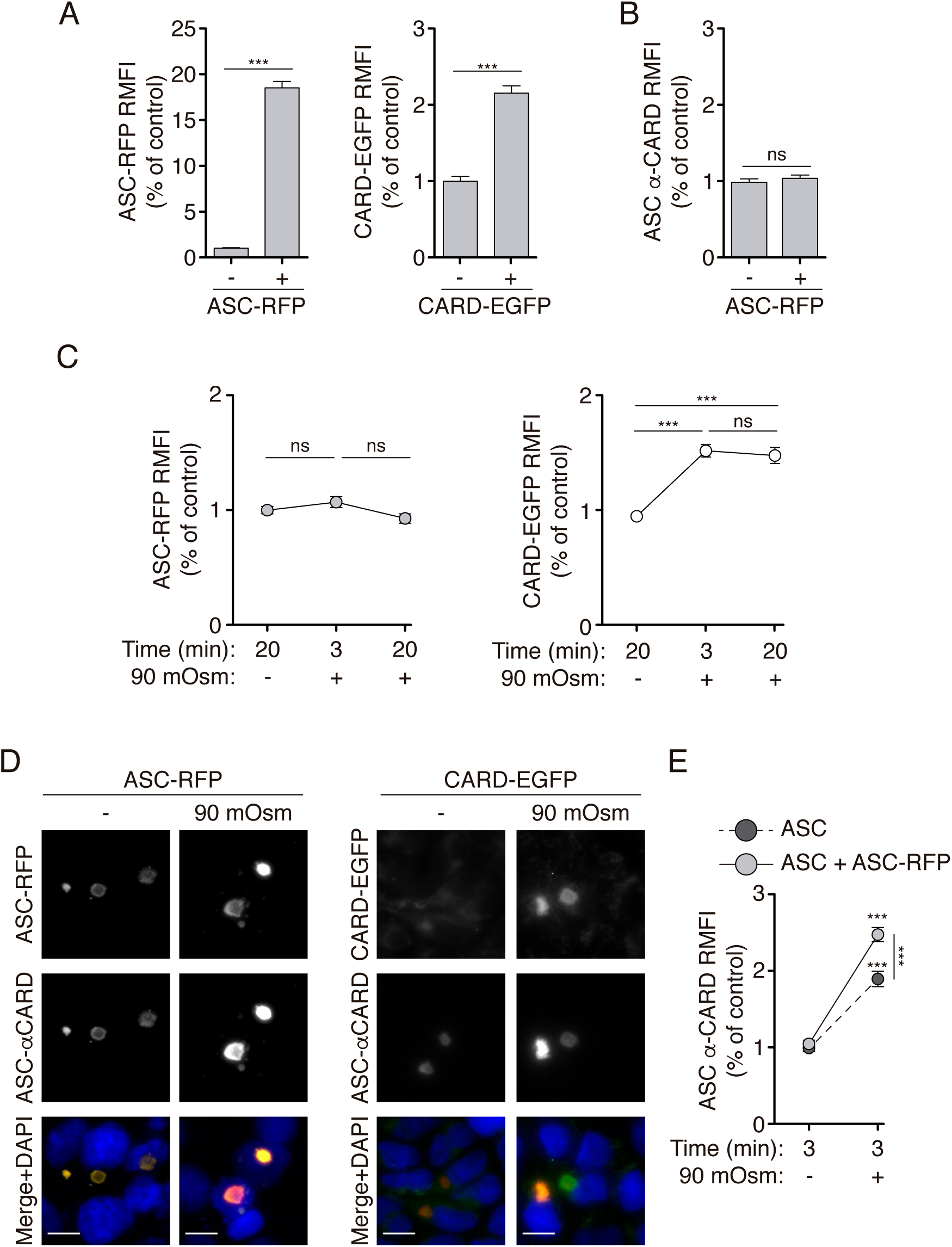
ASC specks recruit more caspase-1^CARD^ domain upon K^+^ efflux. **(A)** Relative mean fluorescence intensity (RMFI) of ASC-RFP (left panel) or CARD_Casp1_-EGFP (right panel) oligomers from HEK293 transfected with untagged ASC (2 μg), then fixed and permeabilized and stained (+) or not (-) with soluble ASC-RFP (left panel) or CARD_Casp1_-EGFP (right panel). Data from *n*= 2-3 independent experiments and quantification of a total of 467 (left panel) and 258 (right panel) ASC-specks. **(B)** RMFI of ASC specks stained with monoclonal α-CARD_ASC_ antibody from HEK293 transfected as in (A) and treated (+) or not (-) with soluble ASC-RFP. Data from *n*= 2-3 independent experiments and quantification of a total of 467 ASC-specks. **(C)** RMFI of ASC specks present in cells transfected as in (A), except that before fixation cells were stimulated (+) or not (-) with hypotonic solution (90 mOsm) for the indicated times. Data from *n*= 2-3 independent experiments and quantification of a total of 464 (left panel) and 586 (right panel) ASC-specks. **(D)** Representative fluorescent photomicrographs of HEK293 cells transfected, treated and stained as in (C) for 20 min. Nuclei were revealed with DAPI (blue); bar 10 μm. **(E)** RMFI of the ASC speck intensity stained with α-CARD_ASC_ from cells transfected as in (A), but before fixation cells were unstimulated (-) or stimulated (+) with hypotonic solution (90 mOsm) for 3 min and then fixed, permeabilized and stained with soluble ASC-RFP (light grey) or nothing (dark grey) and monoclonal α-CARD_ASC_. Data from *n*= 2-3 independent experiments and quantification of a total of 764 ASC-specks.

Finally, to test if this mechanism could have a physiological role, we stimulated the Pyrin inflammasome with the toxin B of *Clostridium difficile* (TcdB) that results in the oligomerization of ASC by Pyrin assembly independently of LPS priming [28]. Caspase-1/11 deficient macrophages were used to avoid pyroptotic cell death upon Pyrin induced ASC oligomerization and no LPS-priming in these experiments assures that NLRP3 inflammasome would not be activated [29]. In this situation, TcdB, but not nigericin or hypotonic stimulation, was able to induce ASC speck formation (**Fig. 6A**). Furthermore, nigericin and hypotonic stimulation were not able to alter the number of specking macrophages upon TcdB stimulation (**Fig. 6A**). However, when nigericin or hypotonicity was applied after TcdB stimulation, the staining intensity for ASC specks determined by immunofluorescence using anti-ASC^CARD^ antibody significantly increased and this increase was blocked using a high extracellular K^+^ solution (**Fig. 6B,C**).

**Figure 6.**
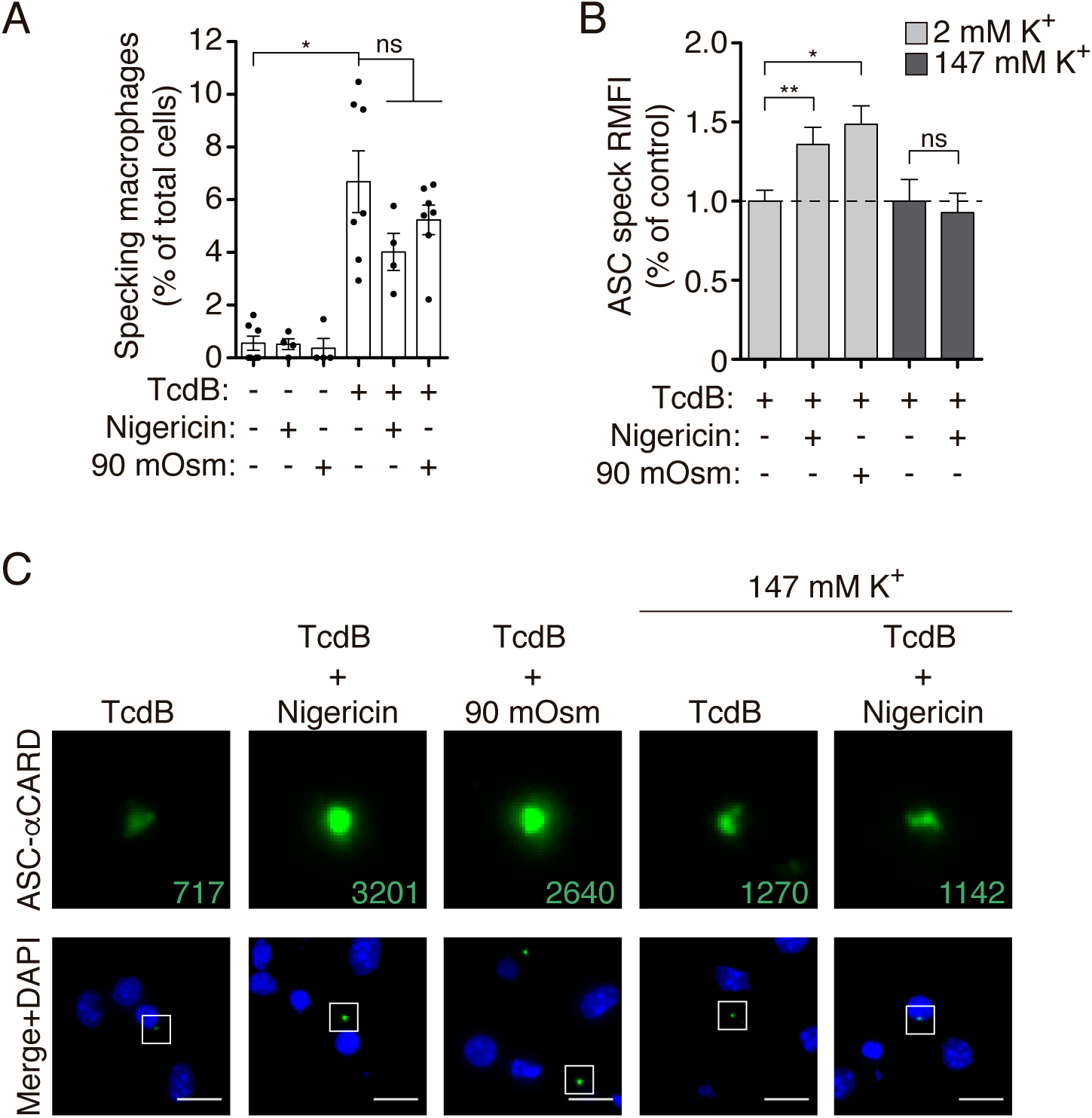
ASC-speck induced by the Pyrin inflammasome present a structural change in response to K^+^ efflux. **(A)** Percentage of ASC specking macrophages derived from *Casp1*^-/-^ mice stimulated with TcdB (1 μg/ml, 1h), nigericin (10 μM, 20 min) or hypotonicity (90 mOsm, 20 min), or nigericin or hypotonicity after TcdB treatment as indicated. ASC was stained with monoclonal α-CARD_ASC_ antibody. Data from *n*= 2 independent experiments and quantification of a total of 1,000 to 2,000 macrophages per treatment. **(B)** Relative mean fluorescence intensity (RMFI) of ASC specks from macrophages treated and stained as in (A), but during treatment a buffer with normal extracellular K^+^ (2 mM K^+^, light gray) or high extracellular K^+^ (147 mM K^+^, dark grey) was used. Data from *n*= 2 independent experiments and quantification of a total of 179 ASC-specks. **(C)** Representative fluorescent micrographs of macrophages treated and stained as in (B). ASC staining with monoclonal α-CARD_ASC_ antibody is amplified for the selected speck in the top panels (numbers denotes RMFI for each of the selected ASC specks) and nuclei were revealed with DAPI (blue); scale bar 10 μm.

In conclusion, our work demonstrates that intracellular K^+^ efflux modulates the ASC oligomer structure at the level of the ASC filament beyond activation of the NLRP3 inflammasome, exposing the ASC^CARD^ domain and favoring the recruitment of caspase-1 into the ASC speck, unrevealing a role for K^+^ efflux favoring caspase-1 activation in non-NLRP3 inflammasomes.

## Discussion

We described in this manuscript that the quaternary structure of the ASC oligomer changes after cellular treatment with the K^+^ ionophore nigericin or during hypotonic induced cell swelling. This structural change can already be observed in ASC filaments assembled by PYD-PYD interactions, and results in a more exposed ASC^CARD^ domain. This allows a better recruitment of the pro-caspase-1^CARD^ domain to the ASC speck.

The NLRP3 inflammasome sensor protein is the only inflammasome component described to be activated by a decrease of intracellular K^+^ and therefore is the only inflammasome triggered by compounds inducing a K^+^ efflux from cells, as specific K^+^-ionophores, activation of ion channels or pore forming toxins [1,2,7]. Also, the activation of caspase-11 induces gasdermin D pyroptotic pores on the plasma membrane that induces an efflux of K^+^ and the downstream activation of the NLRP3 inflammasome [30, 31]. We have found that in conditions of intracellular K^+^ decrease, the ASC speck changes its structure and favor the recruitment of pro-caspase-1, suggesting that initial gasdermin D plasma membrane permeabilization induced by other K^+^-independent inflammasomes could favor the activity of the ASC specks, even if NLRP3 is not present or primed. Therefore intracellular K^+^ concentration appears as a general regulatory mechanism in inflammasome signaling, especially for inflammasomes that are not directly triggered by intracellular K^+^ decrease, as AIM2, NLRC4 or Pyrin [20,21,23,24]. In particular, ASC specks triggered by the activation of the Pyrin inflammasome are also able to change their structure independently of NLRP3 when intracellular K^+^ was decreased.

Recently, we also described that the decrease of intracellular Cl^−^ is important to induce the oligomerization of ASC by NLRP3 activation [32], but under those conditions the resulting ASC oligomers were unable to activate caspase-1, unless cells also present an efflux of K^+^ allowing NLRP3 and NEK7 interaction which in turn activate the inflammasome [32]. The present study shows that the decrease of K^+^ could also modify the ASC speck structure formed by the decrease of intracellular Cl^−^ to favor the recruitment of the pro-caspase-1^CARD^ domain, and therefore could also explain caspase-1 activation in conditions where NLRP3 was activated by Cl^−^ efflux.

The recruitment of ASC by NLRP3 occurs via homotypic PYD-PYD interactions [11, 12] and we have previously described that this interaction is responsive to K^+^ efflux when studied by BRET [33]. Here we found that the oligomeric structure of ASC mainly formed by PYD-PYD interactions is also responsive to a decrease of intracellular K^+^. This idea is further supported with the use of a nanobody that specifically blocks ASC CARD-CARD interactions within the ASC speck that results in filaments of ASC [13]. These ASC filaments with exposed CARD domains are still able to undergo a structural change in response to intracellular K^+^ efflux. The ASC^CARD^ domain is important not only to form the final ASC speck, but also for caspase-1 amplification and signaling [15], therefore the exposure of ASC^CARD^ domains induced by K^+^ efflux could be important to modulate caspase-1 signaling. Since the ASC^PYD^ filaments present a rigid core with a high mobility of the exposed ASC^CARD^ domains [16], our data suggests that this flexibility could be the responsible of the change observed in the ASC oligomer structure induced after cellular K^+^ efflux. The changes in the ASC-ASC BRET signal, suggest overall changes in the quaternary structure of the ASC oligomer, as proximity or orientation of ASC^CARD^ domains change with respect each other. However, we cannot discard that the structural modification in the ASC oligomer observed would be a result of a change in the conformation of individual ASC molecules as regard the orientation of their CARD and PYD domains with respect to each other. The formation of ASC^PYD^ or ASC^CARD^ filaments show that there is no change in the conformation of the individual domains upon oligomerization [11, 34], but these analysis were performed in filaments lacking one of the ASC domains and therefore the orientation of the other domain in the filament is not described, but it is suggested that a flexible linked ASC^CARD^ surround filaments of ASC^PYD^ [11,15,16]. Furthermore, the structural studies of ASC^PYD^ or ASC^CARD^ filaments were resolved in the absence of K^+^ [11,16,34], and our study suggest that the variations on the intracellular ionic environment could be dynamically modulating the structure of the ASC oligomer.

Therefore strategies that could prevent cellular K^+^ efflux could be a general strategy to target different pathologies where the inflammasome is involved [35], as they not only would target NLRP3 activation, but could also reduce the activity of other inflammasomes that signals via ASC oligomerization [20,21,23,24]. Modulation of the structure of the ASC oligomer could be important for different pathologies, as Alzheimer’s disease, where the ASC oligomer is central for the development of the disease [36]. Further understanding of ASC activity regulation in disease will help to refine novel approaches to diseases where the inflammasome is implicated.

## Materials and Methods

### Reagents

*E. coli* LPS O55:B5, DAPI and nigericin were from Sigma-Aldrich; rabbit polyclonal antibody against caspase-1 p10 (M-20), anti-ASC (N-15)-R and horseradish peroxidase-anti-b actin (C4) were from Santa Cruz Biotechnology; mouse monoclonal anti-NLRP3 (Cryo-2) was from Adipogen; mouse monoclonal anti-ASC (TMS-1) was from Biolegend; rabbit polyclonal anti-tubulin and anti-Luciferase from Abcam; ECL horseradish peroxidase conjugated secondary antibody for immunoblot analysis was from GE Healthcare; Alexa Fluor-conjugated donkey IgG secondary antibodies and ProLong Diamond Antifade Mountant with DAPI were from Life technologies; Fluorescence mounting medium was from DAKO; *Clostridium difficile* toxin B (TcdB) was from Enzo.

### Cell culture, treatments and transfection

Bone marrow was obtained from wild type C57BL/6 or *Casp1*^-/-^*Casp11*^-/-^ mice [37] and differentiated to bone marrow derived macrophages (BMDMs) using standard protocols [38]. All animals were maintained under controlled pathogen-free conditions (20 ± 2 °C and 12-hours light-dark cycle), with free access to sterile food and water. HEK293 cells were maintained in DMEM media supplemented with 10% FCS and supplemented with G418 to maintain stable NLRP3-YFP or NLRP3-p.D303N-YFP HEK293 cell lines [19]. Cell lines were routinely confirmed to be free of mycoplasma. Lipofectamine 2000 (Life Technologies) was used for the transfection of HEK293 cells as previously described [39] using different concentrations of plasmids as stated in the figure legends. Cells were stimulated for different times with an isotonic solution (300 mOsm) consisting of (in mM) NaCl 147, HEPES 10, glucose 13, CaCl_2_ 2, MgCl_2_ 1 and KCl 2; hypotonic solution (90 mOsm) was achieved by diluting the solution 1:4 with distilled sterile water.

Alternatively, cells were stimulated with nigericin (10 μM) for different times in isotonic solution.

### ELISA and Western blot

IL-1β release was measured using ELISA kits for mouse IL-1β from R&D following the manufacturer’s instructions. Western blot was carried out as described previously (REF Cam Sebastien). Briefly, protein extracts were resolved in 4–12% polyacrylamide gels and electrotransferred. Membranes were probed with different antibodies for caspase-1, NLRP3, ASC, YFP, luciferase, tubulin or β-actin.

### Bioluminescence resonance energy transfer assay (BRET)

HEK293 cells were co-transfected with a vector encoding for mouse ASC tagged with Luciferase (C-terminus) or YFP (C-terminus) [40]. After 24h, transfected cells were seeded on a poly-L-lysine-coated white 96-well plate the day before to the assay. BRET signal was read after 5 min of coelenterazine-h (5 μM, Invitrogen) addition. Luminescence was detected at 37°C in a Synergy Mx plate reader (Biotek) using two filters for emission at 485 ± 20 nm and 528 ± 20 nm. The BRET ratio was calculated as the difference between the 528 nm and 485 nm emission ratio of R-Luc and YFP-NLRP3 fusion protein and the 530 nm and 485 nm emission ratio of the R-Luc protein alone. Results are expressed in milliBRET (mBRET) units normalized to basal signal as we have previously described [41].

### Immunofluorescence

HEK293 cells or macrophages were seeded on poly-L-lysine coverslips 24 h before use. After transfection and/or stimulation, cells were fixed with 4% formaldehyde, blocked using 2% bovine serum albumin (Sigma-Aldrich) and permeabilized with 0.1% Triton-X100 (Sigma). ASC was stained using a primary antibody anti-ASC (polyclonal anti-PYD ASC, Santa Cruz or monoclonal anti-CARD ASC, Biolegend; 1:1000 dilution) or anti-Luciferase (Abcam; 1:500 dilution) and an Alexa Fluor conjugated secondary antibody (Life technologies; 1:200 dilution). ProLong Diamond Antifade Mountant with DAPI was used as a mounting medium. Images were acquired with a Nikon Eclipse Ti microscope equipped with a 20x S Plan Fluor objective (numerical aperture 0.45), a 40xS Plan Fluor objective (numerical aperture 0.6) and a 60xS Plan Apo Vc objective (numerical aperture 1.40) and a digital Sight DS-QiMc camera (Nikon) with a Z optical spacing of 0.2 μm and 387 nm/447 nm, 482 nm/536 nm, 543 nm/593 nm and 650 nm/668 nm filter sets (Semrock). Images were processed using ImageJ software (NIH) and the maximum-intensity projections images are shown in the results.

### Isolation of soluble caspase-1 CARD-EGFP and ASC-RFP

HEK293 cells transfected with a vector encoding mouse caspase-1 CARD domain tagged with EGFP (0.2 μg) or ASC tagged with RFP (0.2 μg) were lysed in CHAPS (20 mM HEPES-KOH, pH 7.5, 5 mM MgCl2, 0.5 mM EGTA, 0.1 mM PMSF and 0.1% CHAPS) by passing through a syringe on ice (25-gauge needle, 20 times) and the supernatant containing the soluble expressed proteins was obtained by sequential centrifugation at 4°C. The fluorescence in the supernatants was checked using a plate reader and the absence of oligomers was confirmed by fluorescence microscope.

### Soluble caspase-1 CARD-EGFP and ASC-RFP recruitment assay

Poly-L-lysine coverslips seeded HEK293 cells were transfected with mouse ASC (2 μg). 24 h before use. Cells were fixed with 4% formaldehyde after stimulation with nigericin (10 μM) or hypotonic solution (90 mOsm), then washed with PBS, blocked using 2% bovine serum albumin (Sigma) and permeabilized with 0.1% Triton-X100 (Sigma). After that, cells were incubated with soluble caspase-1 CARD domain-EGFP (1:50 dilution in PBS) or ASC-RFP (1:5 dilution in PBS) for 30 min and then fixed for 10 min with 4% formaldehyde, washed with PBS and stained for ASC using a primary antibody against the CARD domain (Biolegend; 1:1000 dilution) followed by an Alexa Fluor conjugated secondary antibody (Life technologies; 1:200 dilution). Nuclei were stained using DAPI (1 μg/ml; 10 min) and the coverslip mounted with fluorescence mounting medium.

### Intensity fluorescence measurement and statistical analysis

The ASC oligomers fluorescence intensity was analyzed using ImageJ software (NIH). For this, a region of interest (ROI) was drawn around the oligomer, measuring the average pixel intensity within each selected area and subtracting an average background value from three selected ROIs within the same image. The fluorescence intensity values obtained were normalized relative to the control inside the same experiment. The data are presented as the mean ± SEM. Data were analyzed using Prism (GraphPad) software using non-parametric tests. Two group comparison was done by the Mann Whitney test, and the Kruskal-Wallis test with Dunn’s multiple-comparison post-test was used to determine the differences among more than two groups (***p<0.001; **p<0.01; *p<0.05; ns, not significant (p>0.05) difference).

## Acknowledgments

The authors thank Dr. Isabelle Couillin (Experimental and Molecular Immunology and Neurogenetics, University of Orleans, Orleans, France) for caspase-1 deficient mice and the SPF-animal house from IMIB-Arrixaca for mice colony maintenance. We thank the members of the Pelegrin’s laboratory for comments and suggestions.

## Author Contributions

F.M-S., V.C., A.T-A., M.C.B. and A.I.G. performed the experiments; F.I.S. provided nanobodies and caspase-1^CARD^ constructions, and interpreted data; F.M-S. and P.P. analyzed and interpreted the data, prepared the figures; F.M-S., V.C. and F.I.S. helped with final manuscript preparation; P.P. conceived the experiments, wrote the paper, provided funding, conception and overall supervision of this study.

## Supplemental material for

**Supplemental Table S1.** Detection of ASC with α-PYD_ASC_ (SantaCruz) and α-CARD_ASC_ (Biolegend) antibodies.

| ASC<br>specie | Tag | Antibody | ASC |  |  |
| --- | --- | --- | --- | --- | --- |
|  |  |  | Soluble | Oligomeric | Oligomeric with<br>low K<sup>+</sup> |
| Human | — | α-PYD<sub>ASC</sub> | + | + | + |
| | | α-CARD<sub>ASC</sub> | + | + | ++ |
| Mouse | — | α-PYD<sub>ASC</sub> | + | + | + |
| | | α-CARD<sub>ASC</sub> | — | + | ++ |
| Mouse | YFP | α-PYD<sub>ASC</sub> | + | + | + |
| | | α-CARD<sub>ASC</sub> | — | — | + |

(-) denotes no staining; (+) denotes staining; (++) denotes an increase of staining.

### Supplemental Figure Legends

**Figure S1.**
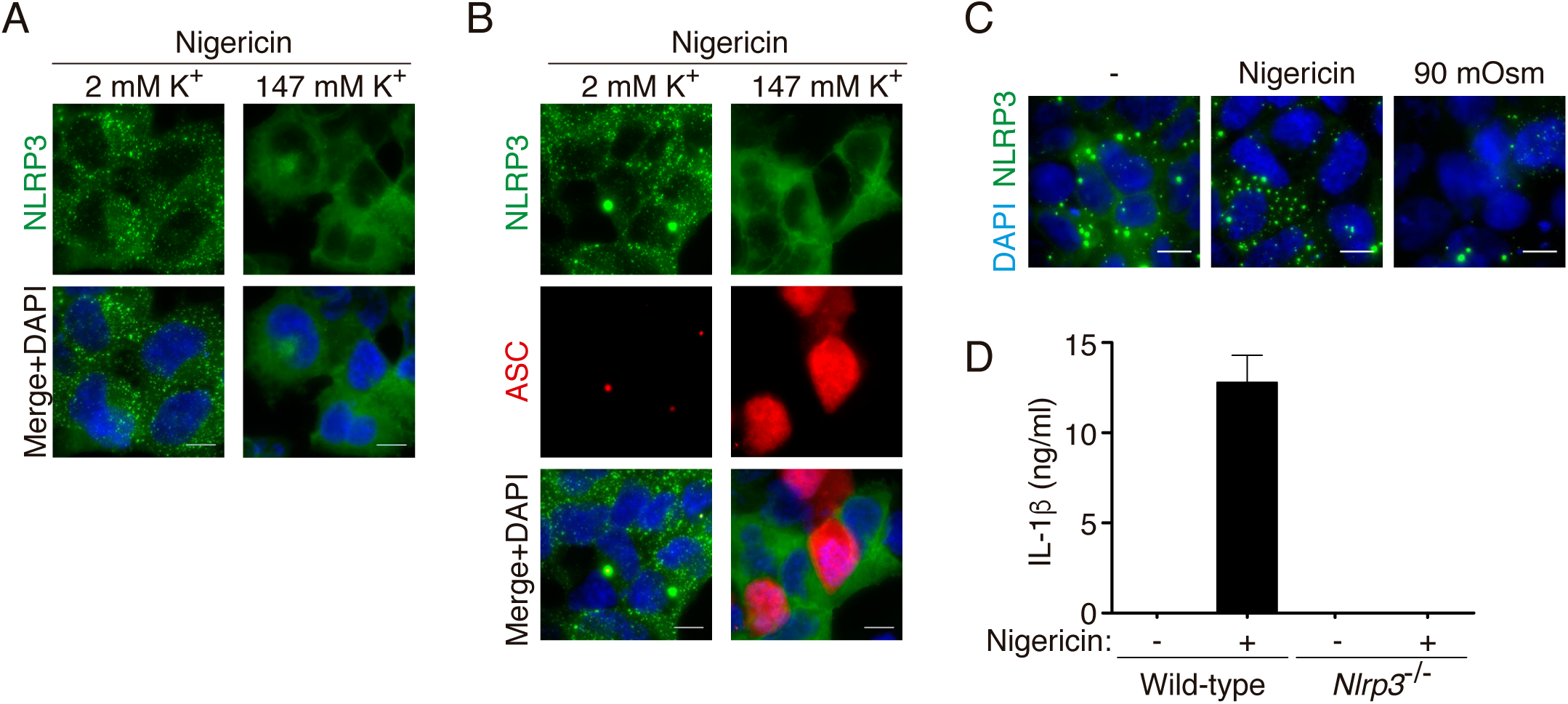
K^+^ efflux induces NLRP3 oligomerization. **(A)** Representative fluorescent micrographs of NLRP3-YFP HEK293 stable cell line and treated with nigericin (10 μM, 30 min) in normal extracellular K^+^ (2 mM K^+^) or high extracellular K^+^ (147 mM K^+^) as indicated. NLRP3-YFP (green) and nuclei was revealed with DAPI (blue); bar 10 μm. **(B)** Representative fluorescent micrographs of NLRP3-YFP HEK293 stable cell line transfected with a plasmid encoding for ASC (1 μg) and treated as in (A). NLRP3-YFP (green), ASC was stained with polyclonal α-PYD_ASC_ antibody (AF647, red) and nuclei was revealed with DAPI (blue); bar 10 μm. **(C)** Representative fluorescent micrographs of NLRP3-YFP D303N HEK293 stable cell line unstimulated (-) or stimulated with nigericin (10 μM, 30 min) or hypotonic solution (90 mOsm, 1 h) as indicated. NLRP3-YFP D303N (green) and nuclei was revealed with DAPI (blue); bar 10 μm. **(D)** ELISA for IL-1β in bone marrow derive macrophage supernatants after LPS (1 µg/ml, 4 h) followed by nigericin (10 μM, 30 min) stimulation.

**Figure S2.**
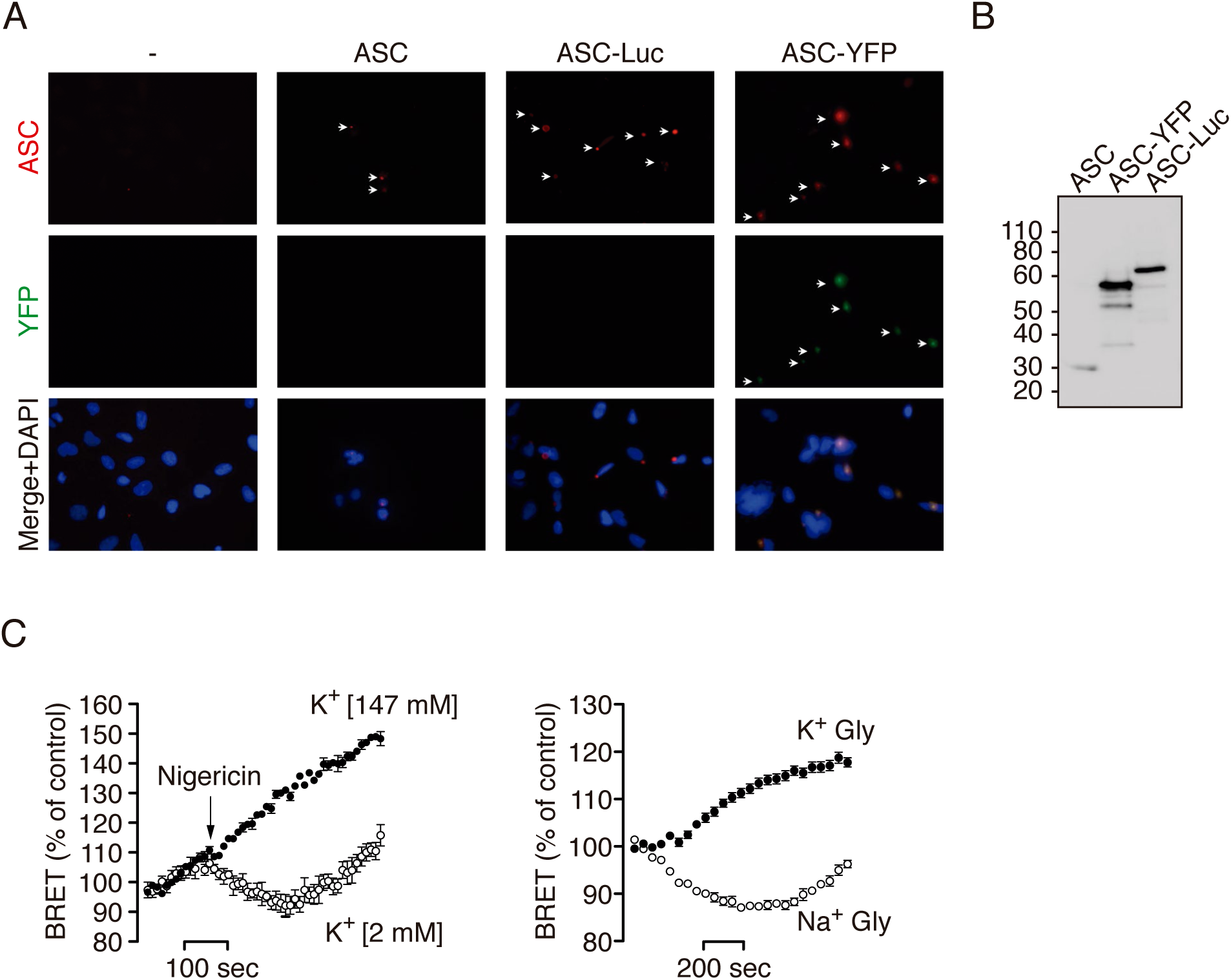
ASC speck conformational change detected by BRET is dependent on K^+^ efflux. **(A)** Representative fluorescent micrographs of HEK293 cells transfected with a plasmid encoding for untagged ASC, ASC-YFP or ASC-Luc. ASC was stained with polyclonal α-PYD_ASC_ antibody (AF647, red), ASC-YFP (green) and nuclei was revealed with DAPI (blue). **(B)** Immunoblot for ASC of cells using the polyclonal α-PYD_ASC_ antibody transfected as in (A). **(C)** Kinetic of net BRET signal in HEK293 cells transfected with the BRET donor ASC-Luc and the acceptor ASC-YFP in response to nigericin (left) or cell swelling induced by glycine solution (Na^+^-Gly, right). Nigericin was applied in normal extracellular K^+^ (2 mM K^+^) or high extracellular K^+^ (147 mM K^+^) buffer as indicated. Cell swelling was performed in sodium (Na^+^-Gly) or potassium (K^+^-Gly) based buffer as indicated.

**Figure S3.**
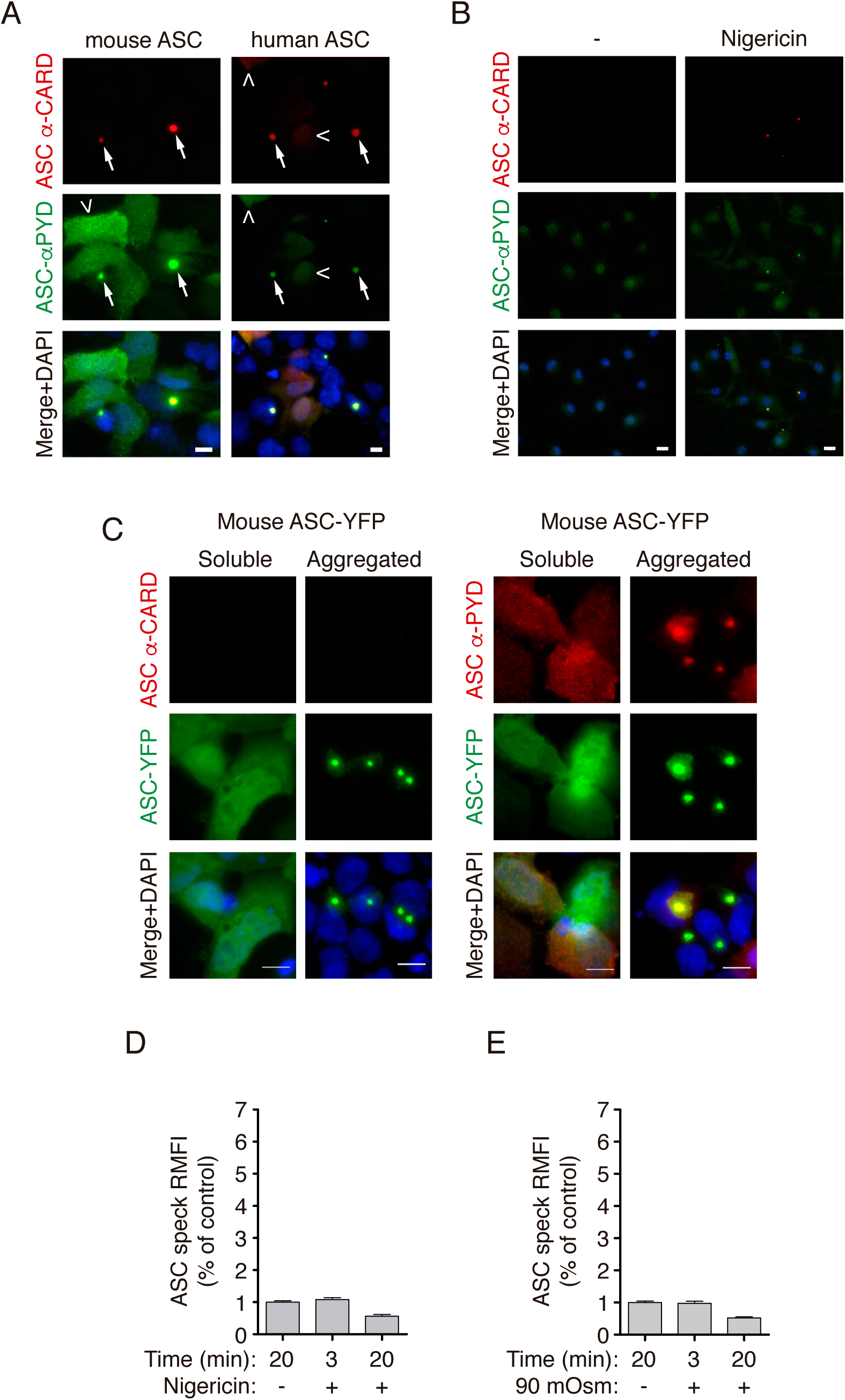
ASC speck is differentially labelled with antibodies. **(A)** Representative fluorescent micrographs of HEK293 cells transfected with a plasmid encoding for untagged mouse ASC (2 μg, left) or untagged human ASC (1 μg, right). ASC was co-stained with monoclonal α-CARD_ASC_ antibody (AF647, red) or with polyclonal α-PYD_ASC_ antibody (AF488, green) and nuclei was revealed with DAPI (blue); bar 10 μm. **(B)** Representative fluorescent micrographs of mouse macrophages primed with LPS (1 μg/ml, 4h) and then stimulated with nigericin (10 μM, 10 min). ASC was stained with monoclonal α-CARD_ASC_ antibody (AF647, red) or with polyclonal α-PYD_ASC_ antibody (AF488, green) and nuclei was revealed with DAPI (blue); bar 10 μm. **(C)** Representative fluorescent micrographs of HEK293 cells transfected with a plasmid encoding for ASC-YFP (0.2 μg). ASC-YFP was staining with monoclonal α-CARD_ASC_ antibody (AF647, red, left panels) or with polyclonal α-PYD_ASC_ antibody (AF647, red, right panels), ASC-YFP (green) and nuclei was revealed with DAPI (blue); bar 10 μm. **(D)** Relative mean fluorescence intensity (RMFI) of ASC-YFP oligomers stained with monoclonal α-CARD_ASC_ antibody from HEK293 transfected with a plasmid encoding for ASC-YFP (0.2 μg), and then unstimulated (-) or stimulated with nigericin (10 μM) for the indicated time in a buffer with high K^+^ (147 mM K^+^). Data from *n*= 2 independent experiments and quantification of a total of 370 ASC-specks. **(E)** RMFI of ASC-YFP oligomers stained with polyclonal α-PYD_ASC_ antibody from HEK293 transfected with a plasmid encoding for ASC-YFP (0.2 μg), and then unstimulated (-) or stimulated with hypotonicity (90 mOsm) for the indicated time. Data from *n*= 2 independent experiments and quantification of a total of 465 ASC-specks.

